# Hepatocyte-Targeted siTAZ Therapy Lowers Liver Fibrosis in NASH Diet-Fed Chimeric Mice with Hepatocyte-Humanized Livers

**DOI:** 10.1101/2023.10.27.564478

**Authors:** Xiaobo Wang, Mary P. Moore, Hongxue Shi, Yoshinari Miyata, Sara K. Donnelly, Daniel R. Radiloff, Ira Tabas

**Author notes:** These authors contributed equally (Xiaobo Wang, Mary P. Moore).

## Abstract

Nonalcoholic steatohepatitis (NASH) is emerging as the most common cause of liver disease. Although many studies in mouse NASH models have suggested therapies, translation to humans is poor, with no approved drugs for NASH. One explanation may lie in inherent differences between mouse and human hepatocytes. We used NASH diet-fed chimeric mice reconstituted with human hepatocytes (hu-liver mice) to test a mechanism-based hepatocyte-targeted siRNA, GalNAc-siTaz, shown previously to block the progression to fibrotic NASH in mice. Mice were reconstituted with human hepatocytes following ablation of endogenous hepatocytes, resulting in ~95% human hepatocyte reconstitution. The mice were then fed a high-fat choline-deficient Lamino acid-defined diet for 6 weeks to induce NASH, followed by six weekly injections of GalNAc-siTAZ to silence hepatocyte-TAZ or control GalNAc-siRNA (GalNAc-control) while still on the NASH diet. The results revealed that GalNAc-siTAZ lowered human hepatic TAZ and IHH, the major TAZ target that promotes liver fibrosis in NASH. Most importantly, GalNAc-siTAZ decreased liver inflammation, hepatocellular injury, hepatic fibrosis, and profibrogenic mediator expression, and profibrotic *NOTCH* vs. GalNAc-control, indicating that GalNAc-siTAZ decreased the progression of NASH in mice reconstituted with human hepatocytes. In conclusion, silencing TAZ in human hepatocytes suppresses liver fibrosis in a hu-liver model of NASH.

**Impact and Implications:** No drugs have yet been approved for NASH, which is a leading cause of liver disease worldwide. The findings here provide support for this therapeutic strategy of using hepatocyte-targeted siTAZ to decrease NASH progression. More generally, the study illustrates how hu-liver NASH mice can be used to evaluate therapeutic hepatocyte-targeted siRNAs to help prioritize future testing in human NASH.

## Introduction

Nonalcoholic fatty liver disease (NAFLD, also known as metabolic dysfunction-associated steatotic liver disease [MASLD]), is characterized by hepatosteatosis (fatty liver), and in ~20-30% of cases, NAFLD progresses to fibrotic non-alcoholic steatohepatitis (NASH, also known as metabolic dysfunction-associated steatohepatitis [MASH]) (1). NASH is emerging as the leading cause of chronic liver disease worldwide, including hepatocellular carcinoma and other comorbidities (1). Disease progression is caused by multiple insults acting on steatotic hepatocytes that result in liver inflammation, hepatocellular death, and, most importantly, liver fibrosis, which correlates best with clinical outcome in NASH (2). No effective therapies currently exist to treat NASH (3), in part due to an incomplete understanding of pro-fibrotic mechanisms in NASH as well as a lack of human-relevant preclinical models of NASH fibrosis, which include mostly mouse models (4).

With regard to mechanisms of NASH fibrosis, we have previously shown that the transcriptional coactivator with PDZ-binding motif (TAZ; encoded by WW domain-containing transcription regulator 1 [WWTR1]) is up-regulated in hepatocytes in mouse and human fibrotic NASH but not steatosis (5, 6). TAZ induces Indian hedgehog (IHH), which activates hepatic stellate cell-derived myofibroblasts (HSCs) and liver fibrosis. Knockdown of hepatocyte TAZ or IHH by experimental genetic methods dampened steatosis-to-fibrotic NASH progression (5, 6). In this context, the last two decades have witnessed the development of therapies that can silence genes specifically in hepatocytes using siRNA conjugated with hepatocyte-specific targeting motifs, notably, N-acetylgalactosamine (GalNAc) (7). Most importantly, several hepatocyte-targeted siRNAs have been recently approved by the FDA for specific disease indications (8-11). Reasoning that GalNAc-siTAZ might have potential as a mechanism-based therapy to suppress the progression to NASH fibrosis, we reported that its administration to mice with early fibrotic NASH suppressed the progression of hepatic inflammation, injury, and fibrosis and lowered the expression of HSC-derived profibrogenic mediators (12).

Since previous studies focused on the effect of GalNAc-siTAZ in mouse hepatocytes, we became interested in a type of chimeric mouse model in which mouse hepatocytes are replaced by human hepatocytes (hu-liver mice) (13, 14). In a recent study, hu-liver mice were fed a NASH-inducing diet and then used to test a systemically administered drug that targets a liver metabolic pathway (15). We reasoned that a mouse NASH model whose livers are populated with human hepatocytes would be particularly valuable in testing hepatocyte-targeted siRNA therapies. Accordingly, we report here on the use of hu-liver NASH mice to investigate the efficacy of GalNAc-siTAZ in preventing the progression of diet-induced NASH fibrosis.

## Materials and methods

### Hu-liver NASH mice

At PhoenixBio (Hiroshima, Japan), cryopreserved human hepatocytes (Lot BD195: 2-year-old, Hispanic female, 1 × 10^6^ hepatocytes, BD Biosciences, Ann Arbor, MI, USA) were transplanted under anesthesia into 2-to 4-week-old hemizygous cDNA-uPA/SCID mice through the spleen. Male hu-liver mice (15 to 18-weeks of age, PXB-mice^®^) were randomized into GalNAc-control and GalNAc-siTAZ groups based on mean body weight and blood human albumin concentration, which was assayed by latex agglutination immunonephelometry (LZ Test ‘Eiken’ U-ALB, Eiken Chemical Co., Ltd., Tokyo, Japan). Based on blood albumin measurements (14), the average replacement index was greater than 90%, which was then verified using anti-STEM121 immunohistochemistry (below). All experiments used a single donor whose hepatocytes were genotyped as C/G at rs738409, indicating they were heterozygous for the PNPLA3 NASH-risk variant (16). The experiments were carried out under procedures and guidelines approved by the Animal Ethics Committee of PhoenixBio, where the in-vivo studies were conducted. The mice were housed in a sterile animal facility with a 12-hour dark/light cycle and free access to food and water. In the first study, the drinking water was supplemented with 0.3 mg/ml ascorbic acid solution (FUJIFILM Wako Pure Chemical Corporation, Osaka, Japan) starting on Day 32 of the study. In the second study, the drinking water was supplemented with ascorbic acid solution for the entire duration of the experiment. NASH was induced by placing the mice on a high-fat choline-deficient L-amino–defined diet (HF-CDAA) (17) (Research Diets, #A06071302, New Jersey, USA) for 6 weeks, followed by weekly subcutaneous injections of GalNAc-siTAZ or GalNAc-Control for another 6 weeks with continued diet feeding. The study was terminated at the end of week 12.

### GalNAc-siRNA

The method of conjugating synthetic triantennary GalNAc ligand targeting the asialoglycoprotein receptor with chemically modified siRNA (18) was applied to Control (non-targeting control) and siTAZ sequences to achieve robust and durable RNAi-mediated TAZ silencing in hepatocytes with the siTAZ conjugate after subcutaneous administration. For Control: Sense 5’-3’: UAUAUCGUACGUACCGUCGUAU, Antisense 5’-3’: AUACGACGGUACGUACGAUdtdt. For siTAZ: Sense 5’-3’: CCUCGAAGCCCUCUUCAACUA, Antisense 5’-3’: UAGUUGAAGAGGGCUUCGAGGAG.

### Histopathologic analysis

Inflammatory cells in H&E-stained liver section images were quantified as the number of mononuclear cells per field (20x objective). For other parameters involving various stains, computerized image analysis (ImageJ) was used to quantify the area stained. The same threshold settings were used for all analyses. For all analyses, we quantified 6 randomly chosen fields per section per mouse. Liver fibrosis was assessed by quantifying Picrosirius (Sirius) red-stained area (Polysciences, #24901). For STEM121 immunohistochemistry, paraffin sections were rehydrated and subjected to antigen retrieval by placing in a pressure cooker for 10 mins in Antigen Unmasking Solution (Vector, H-3300). The slides were then treated with 3% hydrogen peroxide for 10 min and then blocked with Serum-Free Protein Block (Dako, X0909) for 30 min. Sections were incubated overnight with mouse anti-STEM121 primary antibody (Takara, Y40410) and then developed with a DAB substrate kit (Cell Signaling, #8059).

### Immunofluorescence microscopy

Paraffin sections were rehydrated, subjected to antigen retrieval by placing in a pressure cooker for 10 minutes in Antigen Unmasking Solution (Vector, H-3300), and then blocked with serum. Sections were labeled with anti-a-smooth muscle actin (aSMA) (F3777, Sigma), anti-TAZ (Millipore Sigma, HPA007415), anti-Mac2 (Cedarlane, #CL8942AP), anti-Clec4f (R & D, #AF2784), or anti-Ki67 (R & D, #AF7617) antibody overnight, followed by incubation with a fluorophore-conjugated secondary antibody for 1 h. The stained sections were mounted with DAPI-containing mounting medium (Life Technologies, P36935) and then viewed by fluorescence microscopy.

### Plasma transaminase assays

Plasma ALT and AST concentrations were assayed using a kit from TECO Diagnostics (#A526-120) and Biorbyt (orb668836).

### Immunoblotting

Liver protein was extracted using RIPA buffer (Thermo, #89900), and the protein concentration was measured by a BCA assay (Thermo, #23227). Proteins were heated at 100ºC for 5 min, separated by electrophoresis on 4-20% Tris gels (Life technologies, EC60285), and transferred to nitrocellulose membranes (Bio-Rad, #1620115). The membranes were blocked for 30 min at room temperature in Tris-buffered saline and 0.1% Tween 20 (TBST) containing 5% (wt/vol) nonfat milk. The membranes were then incubated in the same buffer with anti-TAZ and anti-β
-actin antibody (#8418, #5125, Cell Signaling) or anti-IHH antibody (#13388-1-AP, Proteintech) at 4°C overnight, using a 1:1000 dilution. The protein bands were detected with horse radish peroxidase-conjugated secondary antibodies and Supersignal West Pico enhanced chemiluminescent solution (Thermo, #34080).

### Quantitative RT-qPCR

Total RNA was extracted from liver tissue using the RNeasy kit (Qiagen, 74106). The quality and concentration of the RNA was assessed by absorbance at 260 and 280 nm using a Thermo Scientific NanoDrop spectrophotometer. cDNA was synthesized from 1 μg total RNA using oligo (dT) and Superscript II (Invitrogen). qPCR was performed with a 7500 Real time PCR system (Applied Biosystems) using SYBR Green Master Mix (Life Technologies, #4367659). The primer sequences are listed in supplementary Table S1.

### Measurement of Hydroxyproline Content of Liver Tissue

The content of hydroxyproline in livers was measured using the Hydroxyproline Colorimetric Assay Kit from Elabscience (E-BC-K061-S). Briefly, liver tissue was hydrolyzed by incubating with the hydrolysis solution at 95°C for 1 hour. After pH adjustment, liver hydrolysates were clarified using activated carbon and then incubated with the reaction reagent for color development. The absorbance was recorded at 550 nm, and hydroxyproline concentrations were determined by comparing them to standards processed simultaneously.

### Statistical Analysis

Data were analyzed using a Student’s t-test for data, with significance set at *p* <0.05.

## Results

Hu-liver mice were placed on the HF-CDAA-NASH diet for 6 weeks, followed by weekly subcutaneous injections with 5 mg/kg GalNAc-siTAZ or GalNAc-control RNA, with continued diet feeding (**Fig. 1A**). The siTAZ used in this study, which targets both mouse and human *WWTR1*, was chosen based on screening experiments in human hepatocytes and mouse NASH models, similar to our previously published strategy for screening GalNAc-siTaz for use in mice (12). The body weights between the two groups were not statistically different (**Fig. 1B, Suppl. Fig.1A**). Immunostaining using an antibody against the human-specific cytoplasmic protein STEM121/SC121 (19) indicated that ~90% of the hepatocytes were of human origin (**Fig. 1C**). At 12 weeks, GalNAc-siTAZ efficiently lowered the mRNA and protein contents of both liver TAZ and its downstream pro-fibrotic mediator IHH (5), with no change in the TAZ-related protein YAP (**Fig. 1D**). GalNAc-siTAZ also lowered the expression of genes related to liver inflammation and hepatic stellate cell activation and profibrotic processes (**Fig. 1E**). Most importantly, while the livers from the GalNAc-control-treated hu-liver NASH mice accumulated inflammatory cells and demonstrated advanced fibrosis, as assessed by H&E and Sirius red staining, respectively, both parameters were decreased by GalNAc-siTAZ (**Fig. 1F**). These changes were accompanied by a 60% decrease in aSMA^+^ cells in the livers (**Fig. 1G**). Human ALT and AST were also lower in the plasma of GalNAc-siTAZ-treated vs. control mice (**Fig. 1H**). In contrast, GalNAc-siTAZ did not significantly affect lipid droplet area in the livers and (**Fig. 1I**). While the reduction in IHH by GalNAc-siTAZ is a likely mechanism for the decrease in liver fibrosis in GalNAc-siTAZ-treated hu-liver NASH mice (5), we considered another possible mechanism. Both pre-clinical and human data suggest an important role for hepatocyte Notch activation in NASH fibrosis (20, 21), and work in other settings has shown that YAP/TAZ can induce *NOTCH* and its ligands (22). We therefore examined the livers for *NOTCH1* and *HES1*, a gene marker of Notch activation. Both *NOTCH1* and *HES1* were substantially lower in the livers of GalNAc-siTAZ-treated vs. control hu-liver NASH mice (**Fig. 1J**), raising the possibility that GalNAc-siTAZ decreases liver fibrosis in NASH by lowering both IHH and Notch signaling.

**Fig. 1.**
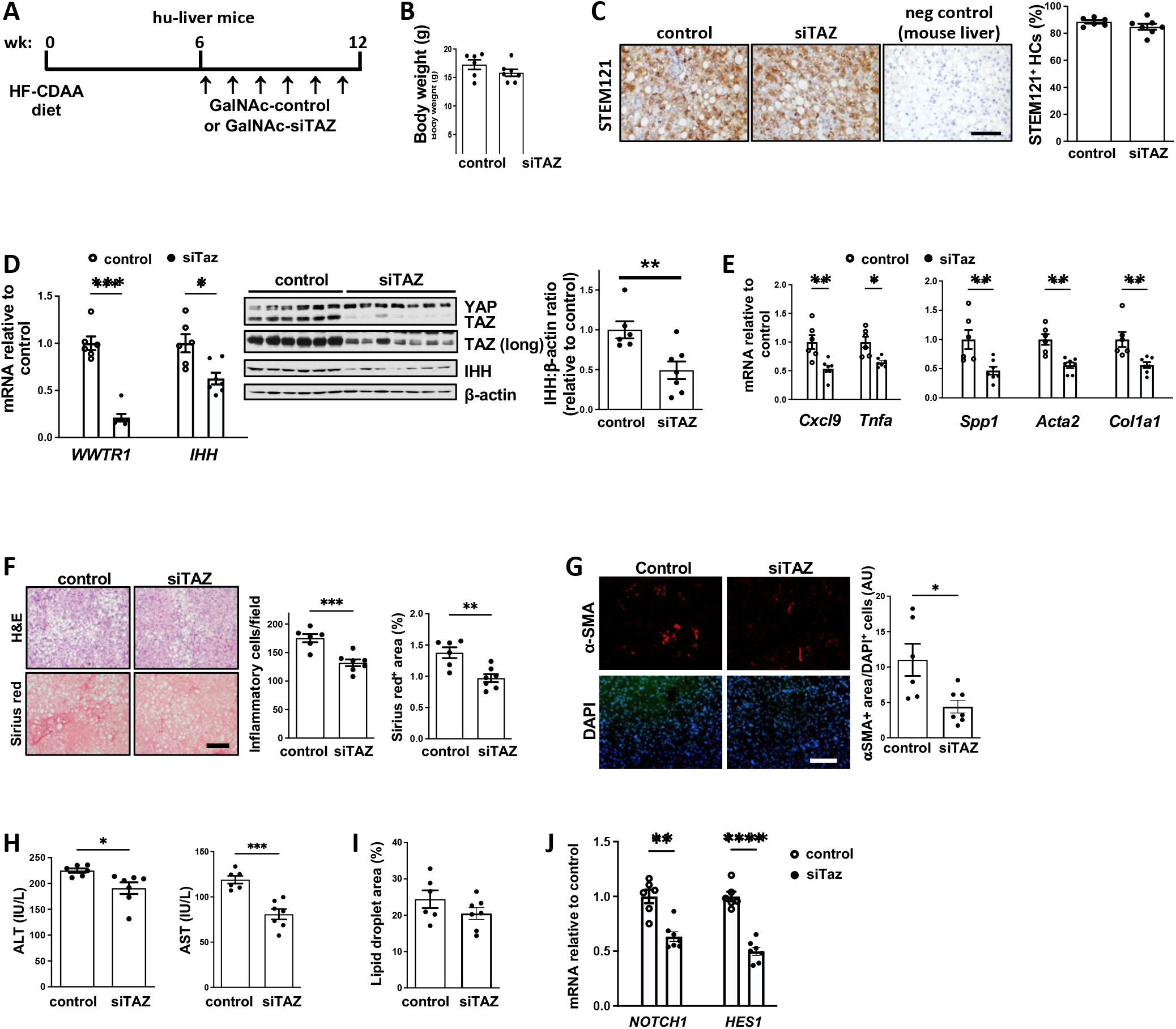
Treatment of hu-liver NASH mice with GalNAc-siTAZ lowers liver inflammation and fibrosis. (A) Experimental scheme. Humanized mice were fed the HF-CDAA diet for 6 weeks and then injected once weekly for 6 additional weeks with 5 mg/kg GalNAc-siTAZ (siTAZ) or control GalNAc construct (control) while still on diet. At 12 weeks, the two cohorts were analyzed for systemic and liver endpoints. (B) Body weight. (C) Livers were immunostained with anti-STEM121, and the percentages of hepatocytes that were STEM121-postive were quantified; liver from a mouse fed the HF-CDAA diet for 12 weeks was used as a negative control for anti-STEM121 staining. Scale bar, 100 μm. (D) Liver *WWTR1* and *IHH* mRNA and immunoblots of YAP, TAZ, IHH, and β-actin; a longer exposure (long) of the TAZ immunoblot is included. The densitometric data for IHH is shown. (E) Livers were assayed for the inflammatory mRNAs *Cxcl9* and *Tnfa* and the fibrosis-related mRNAs *Spp1, Acat2*, and *Col1a1* mRNAs. (F) Livers were stained with H&E (upper row of images) and Sirius red (lower row of images) and then quantified for inflammatory cells per field and percent Sirius red area. Scale bar, 200 μm. (G) Livers were immunostained with anti-α-SMA and quantified for the a-SMA-positive area, which was normalized for cellularity by dividing by the number of DAPI-positive cells. Scale bar, 200 μm. (H) Plasma ALT and AST. (I) Percent lipid droplet area in the livers. (J) Liver *NOTCH1* and *HES1* mRNA. Values shown for all graphs are means □ SEM; n = 6 control mice and 7 siTAZ mice. \**p* <0.05, \*\**p* <0.01, \*\*\**p* <0.001, \*\*\*\**p* <0.0001. The data were analyzed using the Student’s t test.

During this experiment, the mice experienced vitamin C deficiency and some of the mice died before the end of the experiment. Hu-liver mice often develop hypovitaminosis due to their inability to synthesize ascorbic acid (23). To counteract this, the mice are typically provided with a diet containing a small quantity of vitamin C and supplemented with additional vitamins. In our initial study, the vitamin supplement was added to the HF-CDAA diet. However, it was noted that some of the mice avoided the vitamin supplement, possibly due to a shift in dietary preference induced by the HF-CDAA diet. These mice exhibited symptoms consistent with vitamin C deficiency, including low body temperature, reduced activity, and hemorrhage. Upon recognizing these issues, the surviving animals in the study were subsequently provided with vitamin C through their drinking water, commencing on Day 32 of the experiment. Although there was no difference in morbidity or mortality between the control and siTAZ cohorts in this experiment, we conducted a repeat experiment in which vitamin C deficiency was avoided. The human hepatocytes for this experiment came from the same donor as the first experiment. As with the first experiment, the livers contained ~95% human hepatocytes (**Fig. 2A**), and GalNAc-siTAZ efficiently lowered hepatic *WWTR1* and *IHH* (**Fig. 2B**), TAZ-positive human hepatocytes **(Fig. 2C)**, and hepatic inflammation, fibrosis, and plasma ALT, without altering hepatosteatosis and cell proliferation **(Fig. 2D-K)**. In this experiment, although the two cohorts looked to be equally healthy based on overall appearance and activity level, mean body weight of the GalNAc-siTAZ cohort was modestly but significantly lower than that of the control cohort (**Fig. 2L, Suppl. Fig 1B**). We therefore examined a subpopulation of mice in each group whose weights were similar to each other and found that liver fibrosis, as assessed by Sirius red staining, was decreased by GalNAc-siTAZ to a degree similar to that in the full cohort (**Fig. 2M**). Finally, as in the first experiment, the livers of GalNAc-siTAZ-treated hu-liver NASH mice had markedly lower expression of *NOTCH* and *HES1* as well as the Notch downstream effector gene *EPHB2*, which has been implicated in NASH progression (24), compared with control mice (**Fig. 2N**). Thus, in two separate cohorts of hu-liver NASH mice harboring ~95% human hepatocytes, GalNAc-siTAZ dampened the progression to fibrotic NASH.

**Fig. 2.**
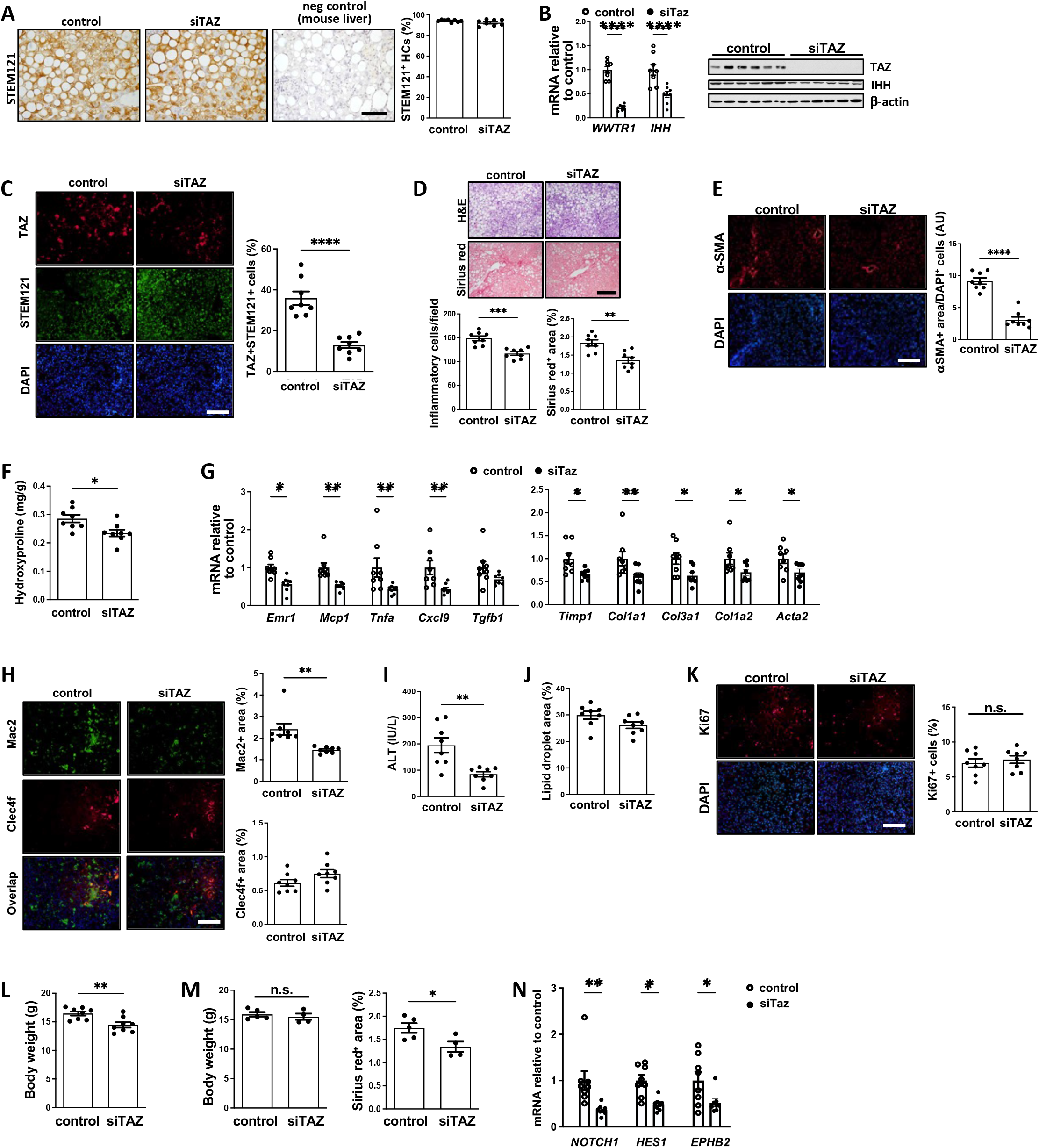
A repeat experiment showing that GalNAc-siTAZ lowers NASH progression in hu-liver mice. A second cohort of hu-liver mice were treated with GalNAc-siTAZ or control GalNAc construct exactly as depicted in Fig. 1A. (A) Livers were immunostained with anti-STEM121, and the percentages of hepatocytes that were STEM121-postive were quantified; liver from a mouse fed the HF-CDAA diet for 12 weeks was used as a negative control for anti-STEM121 staining. Scale bar, 100 □m. (B) Liver *WWTR1* and *IHH* mRNA and immunoblots of TAZ, IHH, and β-actin. (C) Livers were immunostained with anti-TAZ and anti-STEM121 and then quantified for the percent of TAZ-positive cells among STEM121-positive cells. Scale bar, 200 μm. (D) Livers were stained with H&E (upper row of images) and Sirius red (lower row of images) and then quantified for inflammatory cells per field and percent Sirius red area. Scale bar, 200 μm. (E) Livers were immunostained with anti-α-SMA and quantified as in Fig. 1G. Scale bar, 200 μm. (F) Hydroxyproline content. (G) Livers were assayed for the inflammatory mRNAs *Emr1, Mcp1, Tnfa, Cxcl9*, and *Tgfb1* and the fibrosis-related mRNAs *Timp1, Col1a1, Col3a1, Col1a2*, and *Acta2*. (H) Livers were immunostained with anti-Mac2 and anti-Clec4f and quantified for the percent positive area of each. Scale bar, 200 μm. (I) Plasma ALT. (J) Percent lipid droplet area in the livers. (K) Livers were immunostained with anti-Ki67 and quantified for the percent of positive area. Scale bar, 200 μm. (L) Body weight. (M) The livers of a subgroup mice from each cohort (5 control, 4 siTAZ) with similar body weights were assay for percent Sirius red-positive area. (N) Liver *NOTCH1, HES1* and *EPHB2* mRNA. Values shown for all graphs are means ± SEM; n = 8 mice per group. \**p* <0.05, \*\**p* <0.01, \*\*\*\**p* <0.0001. The data were analyzed using the Student’s t test.

## Discussion

Creating therapies to block the progression of fibrotic NASH has proven to be extremely challenging (3), which is particularly alarming given the fact that NASH is emerging as the leading cause of chronic liver disease worldwide (1). The lack of any approved therapies for NASH is likely due to multiple issues, including an incomplete understanding of the mechanisms that promote liver fibrosis in NASH, the possible need for targeting more than one process in NASH (combination therapy), and difficulties in predicting success in humans based on studies in preclinical models (4). The latter problem is due, in part, to inherent differences between hepatocytes in mice, by far the most common species for the initial testing of new therapeutic ideas for NASH, and hepatocytes in humans (25).

To overcome this latter issue, investigators have developed chimeric mice with humanized livers by the transplantation of human hepatocytes into mice whose endogenous hepatocytes have been killed (13, 14). The mice have a hepatocyte gene profile that is 82% similar to hepatocytes isolated from human liver, making it an effective preclinical model to study the potential therapeutic efficacy of novel drugs in vivo (13, 14). Our study was based on a previous report showing that when hu-liver mice are fed a HF-CDAA diet, they display similar histological changes to human NASH, including ballooning, inflammation, apoptosis, regeneration of human hepatocytes, fibrosis, and increases in inflammatory and pro-fibrotic genes in the liver (15). In this report, the authors showed that orally administered elafibranor, a dual PPAR α/δ agonist, improved steatosis and markers of liver injury, but not fibrosis, which is consistent with the results of a previous clinical trial (26). Another study showed that hu-liver mice reconstituted with hepatocytes from a human with a genetic NASH-risk variant, PNPLA3-148M, and then fed a high-fat or Western-type diet had more advanced NASH features compared with high-fat-fed hu-mice reconstituted with non-risk PNPLA3-148I hepatocytes (27).

Advances over the last decade in the area of hepatocyte-targeted siRNA therapy have led to the approval of four drugs for use in humans: patisiran, which targets *TTR* to treat familial amyloid polyneuropathy (8); givosiran, which targets *ALAS1* to treat acute porphyria (9); lumasiran, which targets *HAO1* to treat primary hyperoxaluria type 1 (10); and inclisiran, which targets *PCSK9* to treat hypercholesterolemia (11). Moreover, several other hepatocyte-targeted siRNAs are in clinical trials for a variety of other diseases. We reasoned that for the testing of future hepatocyte-targeted siRNAs, hu-liver NASH mice would help bridge the transition from pure mouse studies to human clinical trials. We chose GalNAc-siTAZ for this purpose given the need for mechanism-based therapies that target proteins involved specifically in NASH liver fibrosis. A combination of in-depth mechanistic studies, analyses of liver tissue and plasma from NASH subjects, molecular-genetic causation studies in NASH mice, and the results of a study testing GalNAc-siTAZ in mouse NASH have all pointed to the promise of this approach (5, 6, 12, 28). Further, we showed that silencing hepatocyte TAZ blocks the progression to hepatocellular carcinoma in NASH mice (29). Accordingly, the data here show that in mice with mostly human hepatocytes, GalNAc-siTAZ blocks fibrotic NASH progression. One mechanism for the GalNAc-siTAZ-induced decreases in fibrosis, inflammation, and liver injury suggested by our previous studies is lowering of the HSC activator IHH (5), and new data here suggest that suppression of the hepatocyte Notch pathway by GalNAc-siTAZ in NASH may be another human-relevant mechanism (20, 21). Previous work has shown that hepatocyte Notch signaling contributes to NASH fibrosis by promoting *Ephb2* expression (24), as knockdown of Ephb2 in hepatocytes ameliorated inflammation and fibrosis in a mouse model of NASH. With regard to inflammation, our data are consistent with GalNAc-siTAZ blocking inflammation in infiltrating monocyte-derived macrophages, which have an inflammatory phenotype in NASH liver (30). Given that only the hepatocytes were of human origin in the mice and that GalNAc-siTAZ targets hepatocytes, not macrophages, the findings suggest that the interaction between human hepatocytes and mouse macrophages remains functional in this model.

In our mouse studies thus far, as in the study herein, we have not observed overt adverse effects of silencing or deleting hepatocyte TAZ, which may be explained in part by the very low expression of TAZ in normal liver and possibly by the ability of the TAZ-related protein YAP to compensate for the loss of TAZ (31). Our study, by design, intervened only after steatosis had developed. However, it will be important in future studies to intervene after early NASH fibrosis has developed, as we did in our mouse NASH study (12). In that study, we subjected mice to the NASH diet for 16 weeks to induce early-stage NASH fibrosis and subsequently administered PBS or GalNAc‐siTaz weekly for an additional 12 weeks while maintaining the mice on the NASH diet. GalNAc‐siTAZ treatment significantly ameliorated both inflammation and fibrosis, and there was also an increase in matrix metalloproteinase activity in liver of the GalNAc‐ siTAZ-treated mice, suggesting that siTAZ may also enhance collagen degradation during NASH regression.

As with all preclinical models, hu-liver NASH mice have limitations, including the SCID background of the mice and the fact that the non-parenchymal liver cells are of mouse origin. The SCID background might affect inflammation in NASH, but the data show that GalNAc-siTAZ reduces the expression of inflammatory genes in a manner that corresponds with a reduction of Mac2^+^ but not resident Clec4f^+^ macrophages, suggesting an effect on infiltrating macrophages (30). These findings indicate that the interaction between human hepatocytes, the site of action of GalNAc-siTAZ, and mouse macrophages remains functional in this model, making it a potentially suitable platform for investigating NASH-related inflammation. Moreover, given a major mechanism by which silencing TAZ in hepatocytes blocks liver fibrosis in NASH (5), i.e., by preventing the induction of the HSC activator IHH, the efficacy of GalNAc-siTAZ in hu-liver NASH mice suggest intact cross-talk between human hepatocytes and mouse HSCs. We conducted this study by using male mice, which are the standard in NASH studies due to male sex being a risk factor for NASH fibrosis in mice and humans (32), and female sex in mice can affect metabolic parameters (33). Nonetheless, future studies in female mice are warranted. Additionally, this study used the HF-CDAA model, but our previous GalNAc-siTaz study used the fructose-palmitate-cholesterol (FPC) diet model, and the results were similar (12). Finally, we think it is noteworthy that the donor human hepatocytes that were used in this study were heterozygous for the potent PNPLA3 NASH-risk variant at rs738409 (16). The success of GalNAc-siTAZ in hu-mice with rs738409-C/G hepatocytes, which is known to exacerbate NASH in hu-liver mice (27), may add further to the evidence suggesting the promise of this therapy. Future studies will be needed to assess GalNAc-siTAZ in hu-mice made from donor hepatocytes without this genetic risk variant.

In conclusion, this study illustrates the potential value of hu-liver mice to test new NASH therapies, especially nucleic acid-based therapies targeting human hepatocytes, with the goal of prioritizing therapeutic candidates for testing in humans. This concept would seem to apply particularly to NASH fibrosis given its rapidly growing prevalence and devastating nature and the absence to date of any approved therapies.

## Abbreviations

GalNAc: N-acetylgalactosamine;
HF-CDAA: high-fat choline-deficient L-amino acid-defined diet;
HSC: hepatic stellate cell; IHH, Indian hedgehog;
NAFLD: non-alcoholic fatty liver disease;
NASH: nonalcoholic steatohepatitis;
SCID: severe combined immunodeficiency.

## Authors’ contributions

XW, MPM, IT, YM, SKD, and DRR were involved in the study conception and experimental design. XW, MPM, and HS conducted liver analyses and plasma and serum assays. XW, MM, and IT drafted the manuscript. All the authors edited the manuscript and approved the submitted version.

## Data sharing statement

Further information and requests for resources, reagents, and data should be directed to the lead contacts, XW and IT.

## Acknowledgments

The authors thank the Takeda Pharmaceutical Company research funding and the Takeda-Drug Discovery Unit Chemistry and Liver Disease teams and the Takeda-Oligochemistry teams for material support and technical input. The Takeda scientists involved in this project included Drs. Bernard Allan, Maria Deato, Jason Pickens, Tony Gibson, Brian Ko, Regina Lemus, Pawan Kumar.

## Figure Legends

**Fig. S1.** Body weight. (A) Body weight of the mice from Figure 1 as a function of weeks on the HF-CDAA diet. (B) Body weight of the mice from Figure 2 as a function of weeks on the HF-CDAA diet.

