## supplementary Figure 1 for "Hepatocyte-Targeted siTAZ Therapy Lowers Liver Fibrosis in NASH Diet-Fed Chimeric Mice with Hepatocyte-Humanized Livers"

**Suppl. Fig.1**

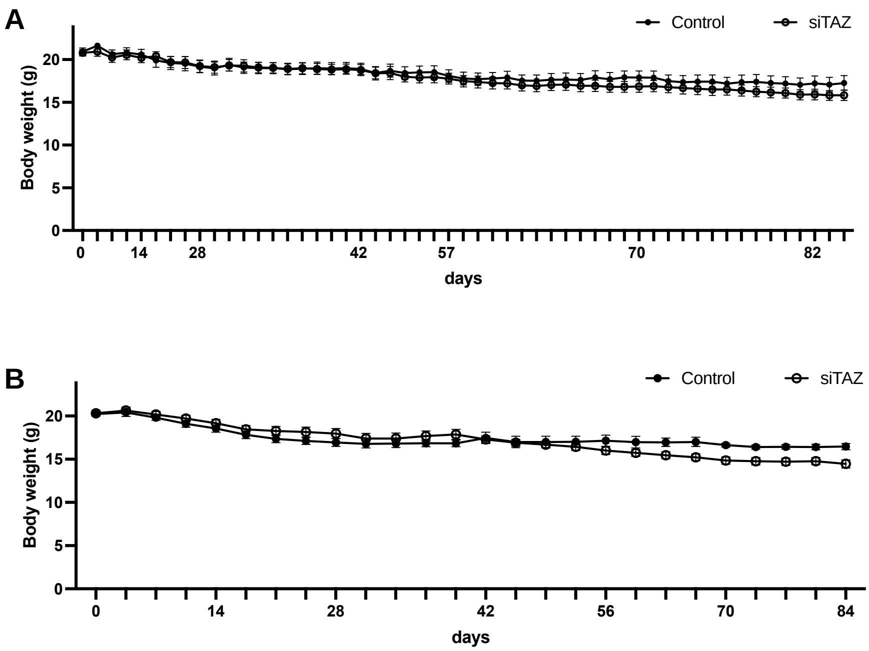
A
Control
siTAZ
Body weight (g)
20
15
10
5
0
0
14
28
42
57
70
82
days
B
Control
siTAZ
Body weight (g)
20
15
10
5
0
0
14
28
42
56
70
84
days

**Table S1 Primers used for QPCR.**

| Primers | Sequences |
|---|---|
| *Hprt* F | TCAGTCAACGGGGGACATAAA |
| *Hprt* R | GGGGCTGTACTGCTTAACCAG |
| *Acta2* F | ATGCTCCCAGGGCTGTTTTCCCAT |
| *Acta2* R | GTGGTGCCAGATCTTTTCCATGTCG |
| *Col1a1* F | GCTCCTCTTAGGGGCCACT |
| *Col1a1* R | CCACGTCTCACCATTGGGG |
| *Col1a2* F | GTAACTTCGTGCCTAGCAACA |
| *Col1a2* R | CCTTTGTCAGAATACTGAGCAGC |
| *Col3a1* F | CTGTAACATGGAAACTGGGGAAA |
| *Col3a1* R | CCATAGCTGAACTGAAAACCACC |
| *Emr1* (*Adgre1*) F | ACCACAATACCTACATGCACC |
| *Emr1* (*Adgre1*) R | AAGCAGGCGAGGAAAAGATAG |
| *Tnfa* F | CTTCTGTCTACTGAACTTCGGG |
| *Tnfa* R | CAGGCTTGTCACTCGAATTTTG |
| *Mcp1* F | TTAAAAACCTGGATCGGAACCAA |
| *Mcp1* R | GCATTAGCTTCAGATTTACGGGT |
| *Spp1* F | CTGACCCATCTCAGAAGCAGAATCT |
| *Spp1* R | TCCATGTGGTCATGGCTTTCATTGG |
| *Timp1* F | CTCAAAGACCTATAGTGCTGGC |
| *Timp1* R | CAAAGTGACGGCTCTGGTAG |
| *Tgfb1 F* | CTCCCGTGGCTTCTAGTGC |
| *Tgfb1 R* | GCCTTAGTTTGGACAGGATCTG |
| *Cxcl9 F* | TCCTTTTGGGCATCATCTTCC |
| *Cxcl9 R* | TTTGTAGTGGATCGTGCCTCG |
| *HPRT F* | CCTGGCGTCGTGATTAGTGAT |
| *HPRT R* | AGACGTTCAGTCCTGTCCATAA |
| *WWTR1 F* | TCCCAGCCAAATCTCGTGATG |
| *WWTR1 R* | AGCGCATTGGGCATACTCAT |
| *IHH F* | AACTCGCTGGCTATCTCGGT |
| *IHH R* | GCCCTCATAATGCAGGGACT |
| *HES1 F* | TCAACACGACACCGGATAAAC |
| *HES1 R* | GCCGCGAGCTATCTTTCTTCA |
| *NOTCH1 F* | GAGGCGTGGCAGACTATGC |
| *NOTCH1 R* | CTTGTACTCCGTCAGCGTGA |
| *EPHB2 F* | CGGCTGCATGTCCCTCATC |
| *EPHB2 R* | GTCCCCGTTACAGTAGAGCTT |

*Hprt*, hypoxanthine guanine phosphoribosyl transferase; *WWTR1*, WW domain containing transcription regulator 1; *Acta2*, α-smooth muscle actin; *Col1a1*, collagen type I alpha 1; *Col1a2*, collagen type I alpha 2; *Col3a1*, collagen, type III, alpha 1; *Emr1* (*Adgre1*), adhesion G protein-coupled receptor E1; *Tnfa*, tumor necrosis factor-α ; *Mcp1*, monocyte chemoattractant protein-1; *Spp1*, osteopontin; *Timp1*, tissue inhibitor of metalloproteinase 1; *Tgfb1,* transforming growth factor beta 1*; Cxcl9,* C-X-C motif chemokine ligand 9; *IHH*, Indian hedgehog; HES1, hes family bHLH transcription factor 1; NOTCH1, notch receptor 1; *EPHB2*, Eph receptor B2.
